# Allo-PED: Leveraging protein language models and structure features for allosteric site prediction

**DOI:** 10.1101/2025.03.28.645953

**Authors:** Xiaochuan Chen, Jianqiang Zheng, Zhenni Huang, Ziqi Xu, Junye Huang, Yanjie Wei, Huiling Zhang

**Affiliations:** College of Mathematics and Information & College of Software Engineering, South China Agricultural University, Guangzhou, 510642, China; College of Materials and Energy, South China Agricultural University, Guangzhou, 510642, China; Shenzhen Institute of Advanced Technology, Chinese Academy of Sciences, Shenzhen, 518055, China

**Keywords:** Allosteric regulation, allosteric site prediction, allosteric pocket prediction, ensemble learning, DCNN

## Abstract

Allosteric regulation plays a pivotal role in modulating protein function and allosteric sites represent a promising target for drug discovery. However, identifying allosteric sites remains challenging due to their structural and evolutionary diversity. Here, we present AlloPED, a novel framework that combines protein language models and machine learning to predict allosteric sites with high accuracy. AlloPED consists of two modules: AlloPED-pocket, an ensemble model leveraging physicochemical features to predict allosteric pockets; and AlloPED-site, a dilated convolutional neural network (DCNN) augmented with a comprehensive attention mechanism for residue-level prediction. AlloPED-pocket achieves state-of-the-art performance on benchmark datasets, yielding an MCC of 0.544 and an AUC of 0.920, outperforming existing methods such as AllositePro and PARS. AlloPED-site further refines predictions using high-dimensional sequence embeddings from the ProtT5 protein language model, achieving a precision of 0.601, a recall of 0.422, and a specificity of 0.661. These results highlight the effectiveness of integrating ensemble learning and deep learning for allosteric site prediction. AlloPED also identifies critical determinants of allosteric sites, including residue clustering coefficients, van der Waals volume, and hydrophobic microenvironments. In summary, this framework provides a robust tool for advancing our understanding of allosteric regulation and facilitating structure-based drug design.

## 1 Introduction

Allosteric regulation is a fundamental biological mechanism in which an effector molecule binds to an allosteric site, inducing conformational and dynamic changes that modulate protein function. This process plays a crucial role in cellular signaling and has been described as “the second greatest secret of life” [1]. Despite its significance, the molecular basis of allosteric regulation remains incompletely understood, and no universal model applies to all proteins[2].

In drug development, allosteric targeting offers advantages over orthosteric approaches. Unlike orthosteric regulators that compete with endogenous ligands, allosteric modulators fine-tune protein activity by enhancing or inhibiting ligand binding at the orthosteric site. They often exhibit greater specificity and fewer side effects [3] since their pharmacological impact plateaus once the allosteric site is saturated. Additionally, allosteric sites experience lower evolutionary pressure than orthosteric sites, making them promising targets for selective drug design with minimal off-target effects. These benefits have fueled growing interest in allosteric drug discovery.

However, identifying allosteric sites remains a major challenge. Most experimentally confirmed sites have been discovered serendipitously, and systematic, efficient prediction methods are still lacking[4]. Computational approaches, including evolutionary analysis, molecular dynamics (MD) simulations, network-based methods, and machine learning, have been explored to address this. Evolutionary analysis identifies conserved residues linked to allosteric regulation [5, 6], while MD simulations assess protein flexibility and conformational shifts under perturbations[7–9]. Network-based approaches analyze residue interaction networks within protein structures [10, 11]. Recently, machine learning has shown promise in allosteric pocket prediction[12–19], though accuracy remains a concern, as current models primarily rely on structural and dynamic features, leaving room for additional predictive factors. More importantly, machine learning-based allosteric site prediction, despite its significance, remains more challenging than pocket prediction and is still in its infancy, necessitating innovative approaches to improve accuracy and expand predictive capabilities.

Protein language models (PLMs) have gained traction in bioinformatics, particularly in functional site prediction. Inspired by natural language processing (NLP), PLMs treat protein sequences as a “biological language,” learning evolutionary relationships, functional patterns, and structural features from large-scale sequence data. This enables powerful functional site prediction. For instance, CLAPE-SMB integrates the ESM-2 pretrained language model with contrastive learning for efficient protein-small molecule binding site prediction [20]. GraphEC employs geometric graph learning with ESMFold for high-accuracy enzyme active site prediction [21], while DPFunc combines sequence data with domain-guided structural features for precise protein function prediction [22]. These studies highlight how PLMs extend beyond sequence analysis, integrating multimodal data and multi-task learning to enhance predictions[23–25].

To develop an efficient and accurate data-driven approach for allosteric site prediction, we propose AlloPED, a novel sequence-based method. AlloPED comprises two components: (1) AlloPED-pocket, an ensemble learning model that predicts allosteric pockets by incorporating structural features; and (2) AlloPED-site, a deep neural network leveraging the ProtT5-XL-UniRef50 (ProtT5)[26] model with 3 billion parameters to generate high-dimensional embeddings. By integrating expanded convolution and attention mechanisms, AlloPED-site captures potential interactions between allosteric sites and other protein regions, enabling precise prediction. Through the combination of ensemble learning and large language model-based deep learning, AlloPED aims to advance allosteric site identification, ultimately benefiting biological research and drug discovery. AlloPED is available at https://github.com/mjcoo/AlloPED

## 2 Materials and Methods

The workflow of AlloPED is systematically outlined in **Fig. 1**. Initially, the ASBench dataset is employed as the training corpus, and protein pockets are identified using the Fpocket algorithm. AlloPED comprises two core modules: Allo-PEDpocket, which employs an ensemble learning architecture to predict allosteric pockets by integrating physicochemical features and residue contact network properties; and Allo-PEDsite, which refines predictions at the residue level using a dilated convolutional neural network augmented with comprehensive attention mechanisms. The final model is validated using an independent test set, ensuring robust generalization performance.

**Fig. 1.**
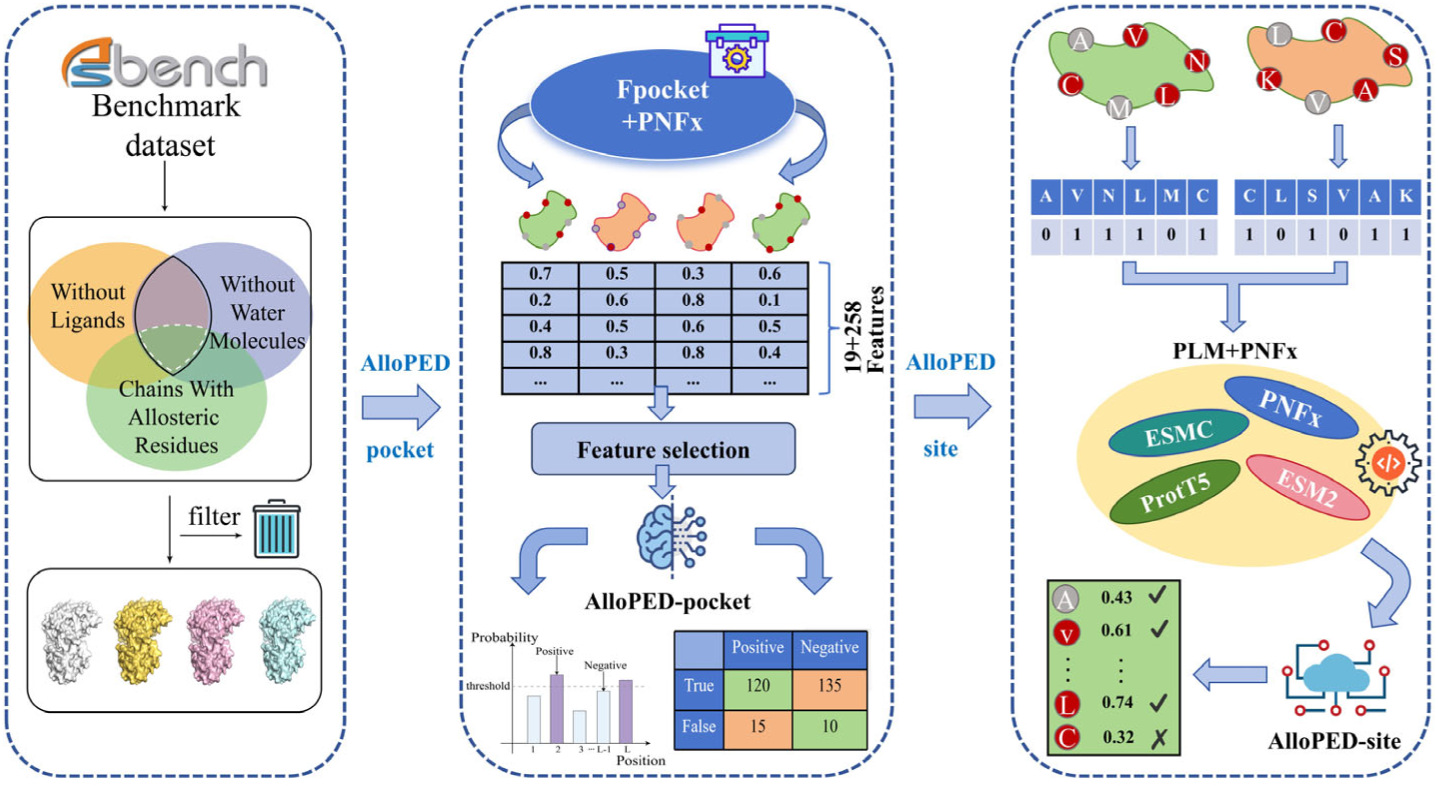
Flowchart of AlloPED for identifying protein allosteric sites.

### 2.1 Data Collection and Preprocessing

The training dataset was derived from ASBench [27], a benchmark collection of allosteric proteins curated from the Allosteric Database (ASD), comprising 146 proteins with X-ray resolution ≤3.0 Å, sequence identity <30%, and experimentally validated allosteric sites. An independent test set of 24 proteins from AllositePro was used for model evaluation. Structural data from PDB files underwent two preprocessing protocols: (1) removal of non-amino acid residues and non-allosteric chains for AlloPED-pocket training, and (2) exclusion of non-amino acid data only for AlloPEDsite training. In this study, Fpocket [28] is used to identify binding pockets on allosteric proteins with default settings. A pocket was labeled positive if ≥25% of its residues overlapped with the true allosteric site; otherwise, it was negative. In the training set (146 proteins), 4,756 pockets were detected (185 positive, 4,571 negative). The test set (24 proteins) contained 53 positive and 745 negative samples. The training dataset included 22776 experiment-validated allosteric sites across 185 positive pockets, while the test set contained 555 sites within 53 positive pockets, which provided sufficient data for comprehensively assessing the model’s performance in allosteric site prediction.

### 2.2 AlloPED-pocket

The workflow of AlloPED-pocket is shown in Fig.2.

**Fig. 2.**
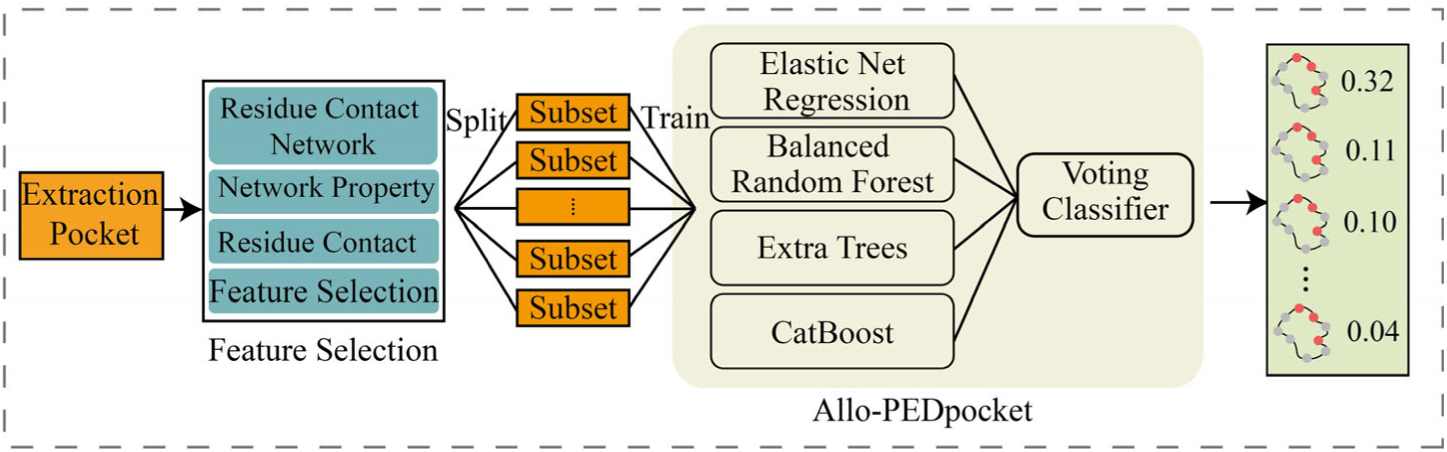
The model architecture of AlloPED-pocket

#### Structure-based features

For allosteric pocket prediction of AlloPED, we use mainly strutural features extracted by Fpocket and PNFx. Fpocket provides 19 physicochemical features as shown in Table S1. PNFx is a feature extraction toolbox developed by our group (https://github.com/mjcoo/PNFx). As shown in Table S2, PNFx provides 258 features such as solvent accsessibility, contact density, clustering coefficient, betweenness centrality, secondary structure and microenvironment properties (Table S2). The detailed description of PNFx can be found at https://github.com/mjcoo/PNFx/feat_list.csv. The Fpocket and PNFx features collectively describe residue and local structural properties of the pocket, thereby enabling systematic analysis of pocket-level functional determinants.

#### Feature Selection

We implemented a two-stage feature optimization strategy to balance dimensionality and model performance. In the first stage, the Minimum Redundancy Maximum Relevance (mRMR) algorithm filters the 258-dimensional structural feature space, using a correlation threshold of 15 to maximize relevance while minimizing redundancy, enhancing computational efficiency and interpretability. Next, pocket-level feature descriptors are constructed by aggregating mean, minimum, and maximum values of amino acid features within each binding pocket. In the second stage, Recursive Feature Elimination with Cross-Validation (RFECV) with a random forest model refines feature selection through iterative backward elimination, dynamically assessing each feature’s contribution to model performance.

#### Allosteric Pocket Prediction Model

AlloPED-pocket utilizes a Soft Voting Ensemble Learning model, combining CatBoost, Elastic Net Regression, Balanced Random Forest, and Extra Trees to improve prediction accuracy. CatBoost refines weak learners within a Boosting framework, excelling in handling high-dimensional data. Elastic Net integrates L1 and L2 regularization, making it suitable for correlated features. Balanced Random Forest mitigates class imbalance through resampling, while Extra Trees enhances generalization via randomized node splitting.

For training, the dataset was divided into 10 balanced groups, ensuring each positive sample was paired with a randomly selected negative sample sharing the same PDB ID. GroupKFold with 5-fold cross-validation trained the ensemble model and generated prediction probabilities for each base learner. A dynamic thresholding approach was applied, with Youden’s index optimizing the decision threshold via ROC curve analysis:

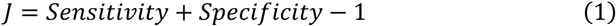

Instead of using a fixed 0.5 threshold, this method ensured a better balance between recall and specificity.

### 2.3 AlloPED-site

The workflow of AlloPED-site is shown in Fig.3.

**Fig. 3.**
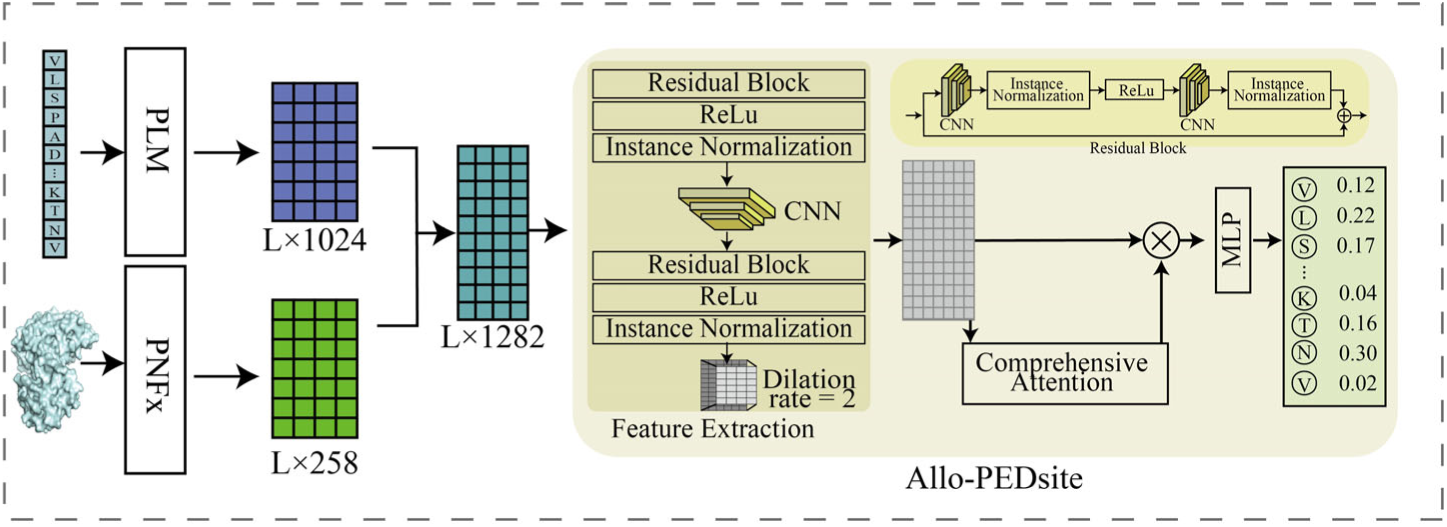
The model architecture of AlloPED-site

#### Sequence and structural features

AlloPED-site integrates sequence embeddings from PLM and structural features from PNFx to capture evolutionary and functional patterns. In our preliminary work, we compared several PLMs (ESM2, ESMC-600M, PROtt5-3B). Since ProtT5-3B performed best, AlloPED-site chose it as the protein language model for sequence encoding. Structural features from PNFx, identical to those used in AlloPED-pocket, include 258 physicochemical and topological descriptors. These multimodal features are concatenated into a high-dimensional vector and processed using a sliding window approach. Unlike AlloPED-pocket’s two-stage mRMR-RFECV feature selection, AlloPED-site employs end-to-end neural architecture to automatically select discriminative features, enabling adaptive learning of sequence-structure relationships for allosteric site prediction.

#### Allosteric Site Prediction Model

Before training AlloPED-site, protein structural data underwent preprocessing, including feature vector extraction, normalization, and window-based sampling to enhance local feature capture. The model, based on a Dilated Convolutional Neural Network (DCNN), incorporates dilated convolution, residual connections, mixed attention mechanisms, and focal loss functions to improve allosteric site prediction. Dilated convolution expands the receptive field to capture long-range dependencies, while residual connections mitigate gradient vanishing and enhance feature propagation. Mixed attention mechanisms prioritize critical feature regions, and the focal loss function emphasizes hard-to-classify samples to address class imbalance.

Training employed Kaiming normal initialization for convolutional weights and the Adam optimizer with a cyclical learning rate, following 5-fold cross-validation. Data was split into five subsets, with four for training and one for validation per fold, allowing hyperparameter tuning and refinement. The final model’s performance was evaluated across all folds, ensuring reliability in allosteric site prediction.

#### Feature Extraction Module

Each protein’s feature matrix X_inuput_ is first mapped to a 512-dimensional space through an initial convolutional layer *X*_1_ = *Conv*1*d*(*X_inuput_*), followed by instance normalization *X*_2_ = *InstanceNorm*1*d*(*X*_1_) and ReLU activation *X*_3_ = *ReLU*(*X*_2_) for stability and non-linearity. A residual block *X*_4_ = *ResidualBlock*(*X*_3_,512), another convolutional layer *X*_5_ = *Conv*1*d*(*X*_4_; *dilation* = 2), and instance normalization *X*_5_ = *InstanceNorm*1*d*(*X*_5_) refine the features before passing through a second residual block. Dilated convolution expands the receptive field, capturing long-range dependencies and broader contextual information. Finally, the feature extraction module outputs *X*_6_ = *ResidualBlock*(*X*_5_,512).

#### ResidualBlock

The residual block consists of two convolutional layers with instance normalization and ReLU activation 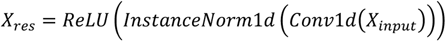. The shortcut connection mitigates gradient vanishing, promoting feature reuse. To enhance representation, channel attention is applied, where X_inuput_ undergoes global average pooling and instance normalization to generate attention weights 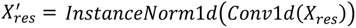. These weights recalibrate the output 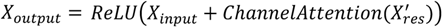, emphasizing key feature channels.

#### ChannelAttension

Channel attention highlights important feature channels by reducing dimensions to 16. By learning inter-channel relationships, the model selects the most valuable features for each position.

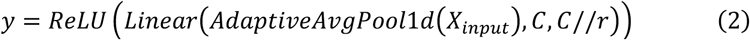

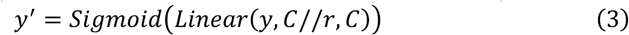

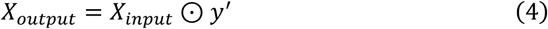

#### Comprehensive Attention Module

This module enhances feature representation by integrating spatial and channel attention. The input matrix X_6_ undergoes convolution *A_spatial_* = *Conv*1*d*(*X*_6_) and Sigmoid activation to generate spatial attention weights 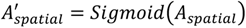, highlighting key regions for allosteric site prediction. Simultaneously, channel attention produces *y_attn_* = *ChannelAttention*(*X*_6_). The final attention weight 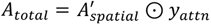, obtained by multiplying both, is applied to *X*_6_, resulting in 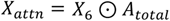, which enhances focus on crucial regions and channels, improving prediction accuracy.

#### Classification Module

This module maps weighted feature vectors to allosteric site labels through binary classification. The weighted feature map X_attn_ is transposed for fully connected layer operations, followed by dimensionality reduction and feature extraction via fully connected layers and layer normalization. ReLU activation enhances non-linearity, while Dropout prevents overfitting. The final classification layer applies a Sigmoid function to map output values to [0,1].

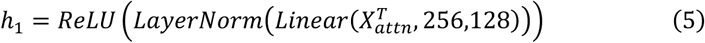

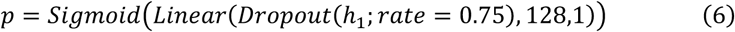

where p represents the probability of an allosteric site

#### Loss Function

The model uses the Focal Loss function as its optimization objective to effectively address class imbalance issues and increase the model’s attention to hard-to-classify samples.

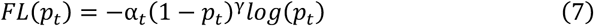

where p_t_ represents the predicted probability of belonging to the positive class. The class imbalance coefficient α_t_ is set to 0.85, and the hard sample focusing parameter γ is set to 4.0, to increase the attention paid to positive examples with smaller quantities.

### 2.4 Performance Evaluation Metrics

To assess the performance of AlloPED, we conducted a thorough evaluation using an independent test dataset. The test dataset underwent the same feature extraction and preprocessing pipeline as the training data to ensure consistency in evaluation. To quantify the model’s predictive performance, we used several standard classification metrics: Precision (PRE), Specificity (SPE), Recall, and Matthews Correlation Coefficient (MCC). These metrics are defined as follows:

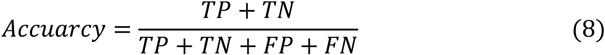

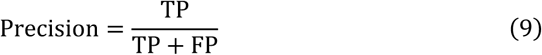

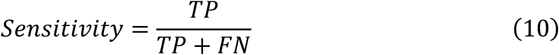

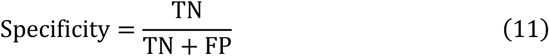

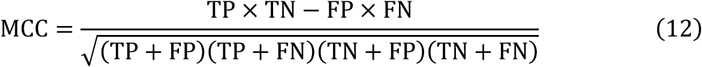

where TP, TN, FP, and FN represent the number of true positives, true negatives, false positives, and false negatives, respectively. These metrics collectively provide a comprehensive evaluation of the model’s performance. We also utilized the Area Under the Receiver Operating Characteristic Curve (AUC) to assess the model’s ability to distinguish between positive and negative classes, offering an additional measure of classification accuracy.

## 3 Results

### 3.1 Characteristic analysis of allosteric sites

Allosteric sites exhibit distinct evolutionary and structural characteristics compared to functional sites. Evolutionarily, they experience reduced selective pressure, enabling adaptive mutations (e.g., asparagine-related features in RNR and Asp mutations in KRAS), while maintaining functional conservation through conformational plasticity. Structurally, allosteric sites are smaller, more dynamic, and enriched in hydrophobic residues, with shallow binding pockets and network connectivity critical for signal propagation. These properties, including high clustering coefficients and β-sheet structures in hinge regions, align with previous findings highlighting their roles in regulatory flexibility and pathological processes.

In this study, we analyzed the physicochemical properties strongly associated with allosteric sites. In the first stage of feature selection, we applied mRMR analysis to systematically screen allosteric site features and compare them with other functional sites (**Fig. 4** and **Fig. 5**). The allosteric score and orthosteric score are relevance values calculated through mRMR that indicate how strongly each feature correlates with allosteric or orthosteric sites, respectively. Higher scores indicate that the feature is more important in predicting that specific type of site. The results indicated that allosteric sites exhibit low sequence specificity, as seen in their conservation scores and hydrophobic microenvironment. Large-scale sequence alignment revealed an average conservation score of 0.63 for allosteric sites, significantly lower than orthosteric (active) sites (0.83, P = 1.26 × 10^-23^) [29], suggesting greater sequence plasticity during evolution. Additionally, allosteric sites are enriched in hydrophobic residues such as isoleucine (Ile), whereas active sites favor rigid residues like tryptophan (Trp) and phenylalanine (Phe) [29, 30]. Statistical analysis of 2,937 proteins (69401 allosteric sites in ASD database) further supported this trend (**Fig. 6**), with isoleucine appearing 4,317 times in allosteric sites (6.22%). Moreover, differences in negative charge distribution suggest that the electrostatic microenvironment of allosteric sites may dynamically regulate ligand binding, as observed in the HisH-HisF system [31].

**Fig. 4.**
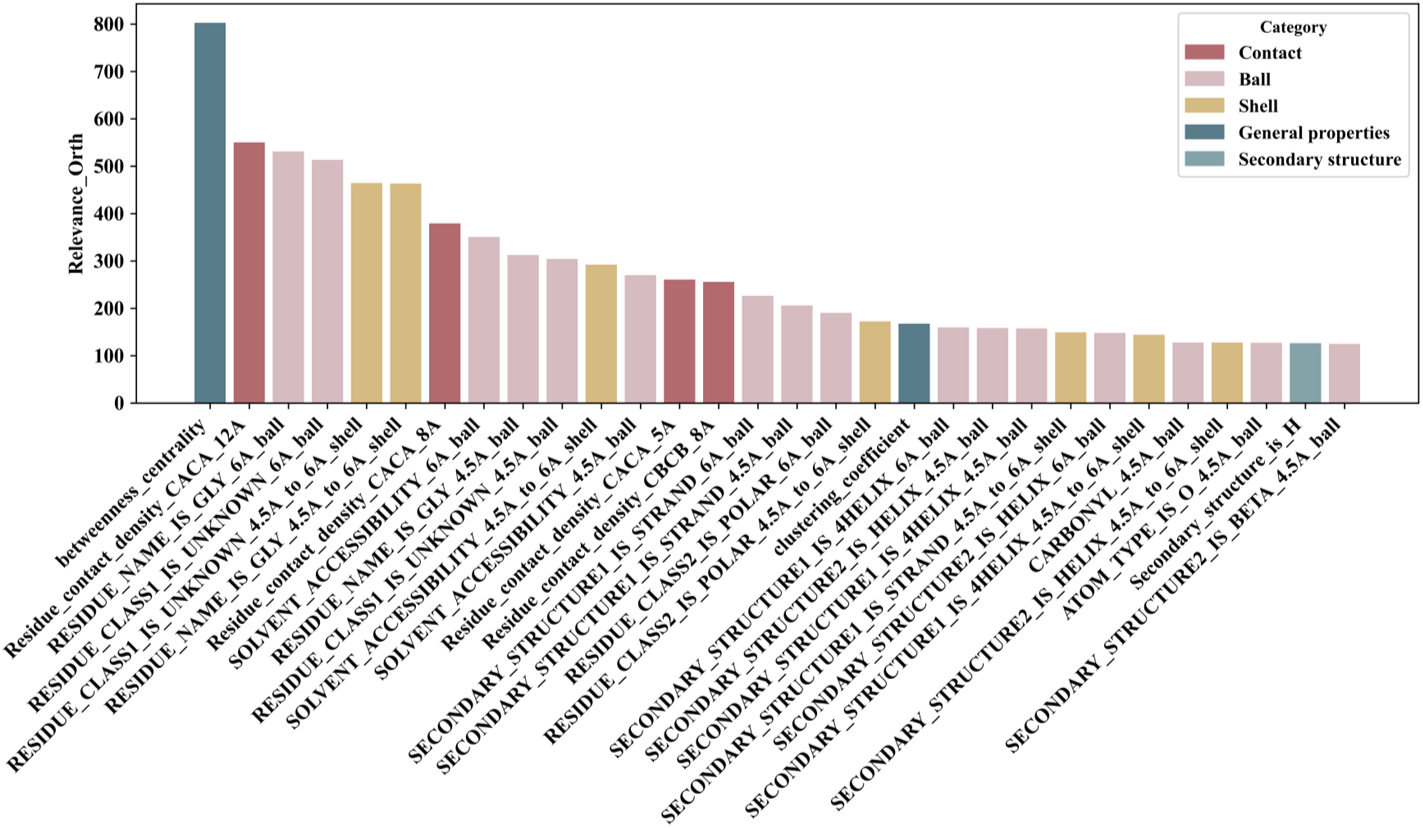
Top 30 features ranked by orthoteric score using mRMR analysis.

**Fig. 5.**
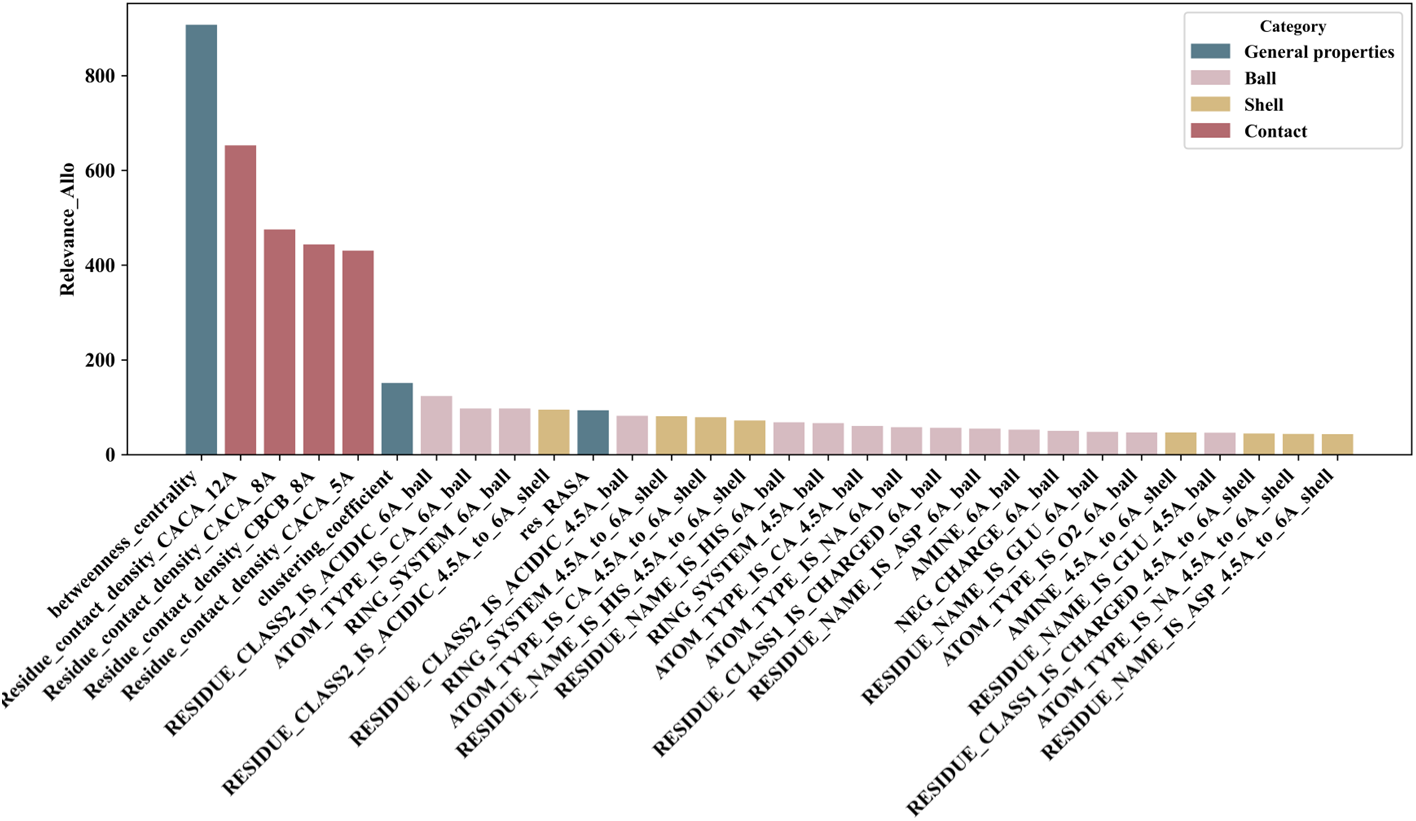
Top 30 features ranked by allosteric score using mRMR analysis.

**Fig. 6.**
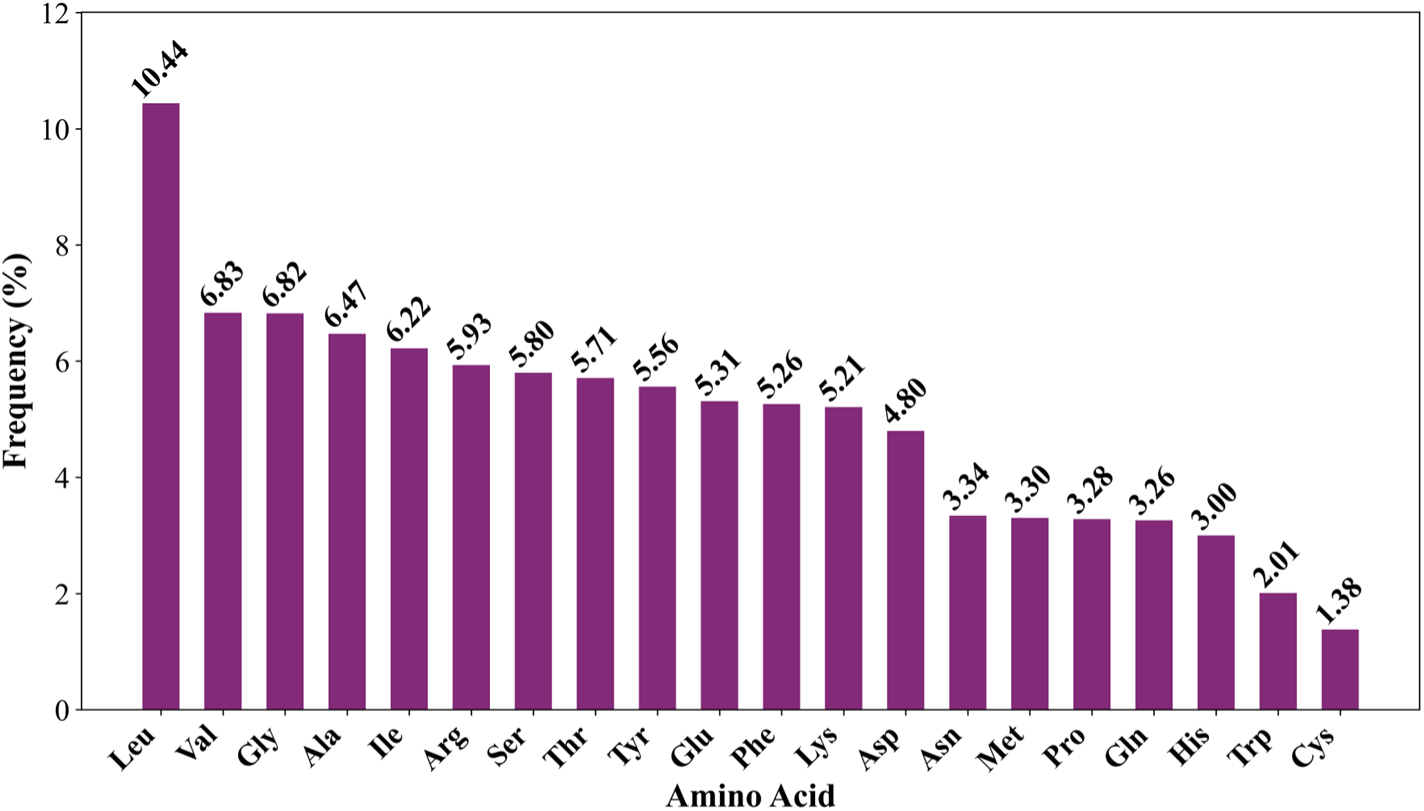
Frequency distribution of amino acids at allosteric sites according to 2,937 allosteric proteins.

### 3.2 Performance of AlloPED-pocket

Feature selection plays a crucial role in improving AlloPED-pocket’s predictive performance. Using mRMR analysis, we identified the most informative features, enhancing accuracy and generalization. Clustering coefficient and van der Waals volume emerged as key predictors, as they are not only statistically linked to allosteric sites but also biologically relevant—clustering coefficient reflects their role in regulatory networks, while van der Waals volume captures their compact structural properties. Additionally, mRMR eliminated redundant features, ensuring the model focuses on the most relevant data. To refine feature selection, we computed aggregated statistics (e.g., mean, minimum, and maximum values) for amino acid properties to construct pocket-level descriptors. Finally, RFECV identified variables that best characterize allosteric pockets, optimizing AlloPED-pocket’s predictive performance.

Table 1 presents the comparison results of AlloPED-pocket with AllositePro, Allosite, PARS, and other methods. As shown in **Table 1**, our model achieved a Matthews correlation coefficient (MCC) of 0.544, significantly outperforming other methods, with a 40.2% improvement over AllositePro. This indicates that our model has certain advantages in addressing the issue of class imbalance. Additionally, our model’s precision was 47.1%, which represents a significant improvement over AllositePro (33.3%), Allosite (24.5%), and PARS (18.0%). Notably, our model achieved the best performance in terms of specificity and AUC, demonstrating its superior ability to avoid false positives among negative samples and indicating stronger generalization capabilities compared to other methods.

**Table 1.** Performance comparison of AlloPED with other methods on the test set.

| Methods | ACC | SPE | PRE | Recall | MCC | AUC |
| --- | --- | --- | --- | --- | --- | --- |
| PARS | 0.708 | 0.756 | 0.180 | 0.375 | 0.099 | - |
| Allosite | 0.828 | 0.858 | 0.245 | 0.500 | 0.264 | - |
| AllositePro | 0.863 | 0.885 | 0.333 | 0.625 | 0.388 | - |
| AlloPED-pocket | 0.952 | 0.965 | 0.471 | 0.686 | 0.544 | 0.920 |

### 3.3 Performance of AlloPED-site

During model training, we concatenated the features generated by ProtT5 with 258 structural features. This approach, leveraging the structural characteristics of AlloPED-site’s deep learning model, helps avoid missing certain potential features that could determine allosteric sites in previous feature selection processes. By training and evaluating the AlloPED-site model using different window sizes (ranging from 5 to 45 with an interval of 4), we systematically calculated the model’s performance across various window sizes (**Fig.7**). The experimental results showed a significant trend in the performance metrics as the window size increased. The model achieved optimal AUC (0.72) and AUPRC (0.67) when the window size was 5, with precision and recall values of 0.63 and 0.56, respectively, indicating a good balance. As the window size increased, we observed a noticeable decline in the AUPRC metric, while the AUC showed only a slight decrease. Particularly in terms of precision and recall, both metrics exhibited a downward trend, with recall experiencing a significant drop. The F1 score also reached its highest value (0.59) at a window size of 5, while the MCC metric remained relatively stable across different window sizes but performed better at smaller windows.

**Fig. 7.**
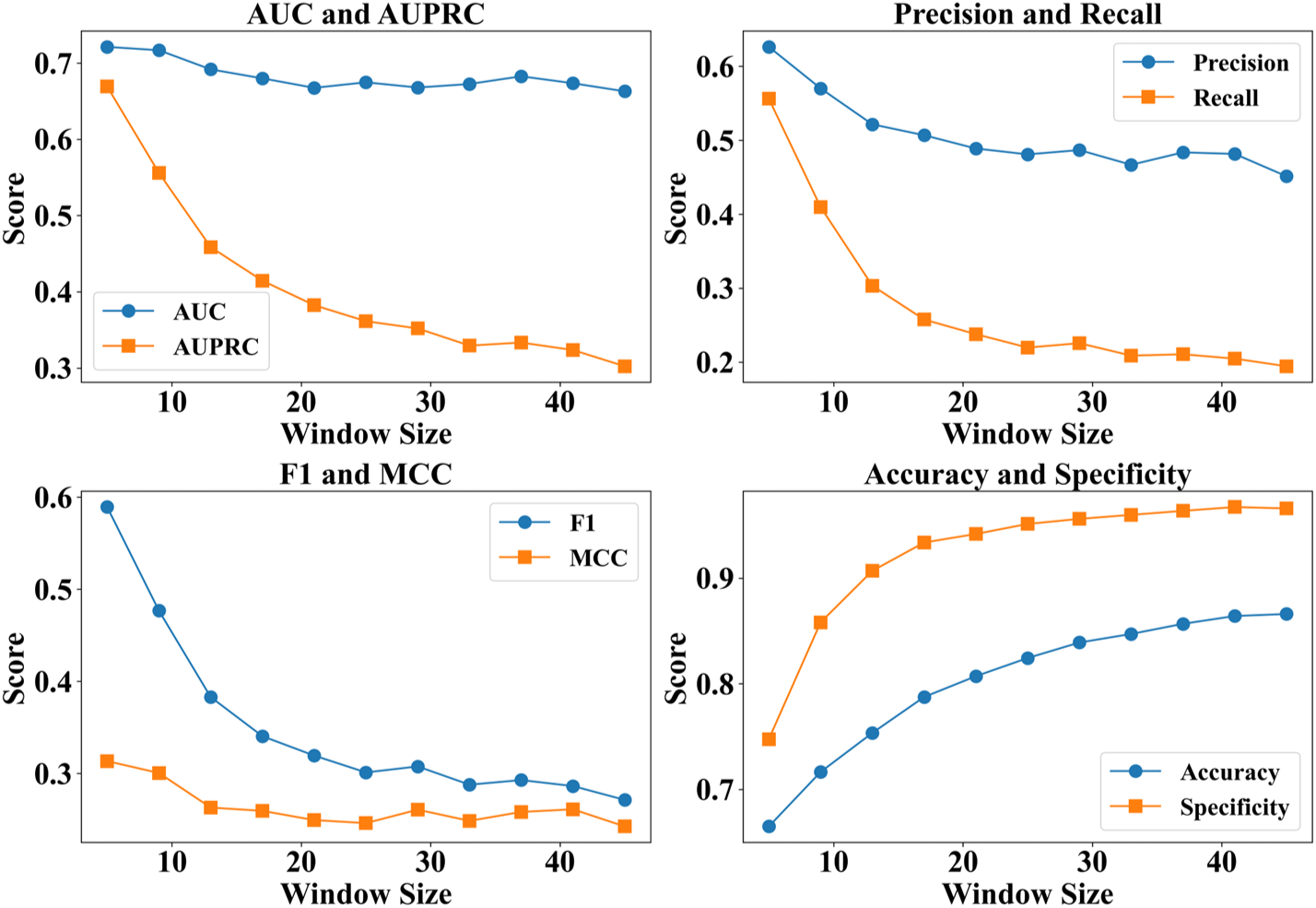
Performance of AlloPED-site across different window sizes

**Fig. 8.**
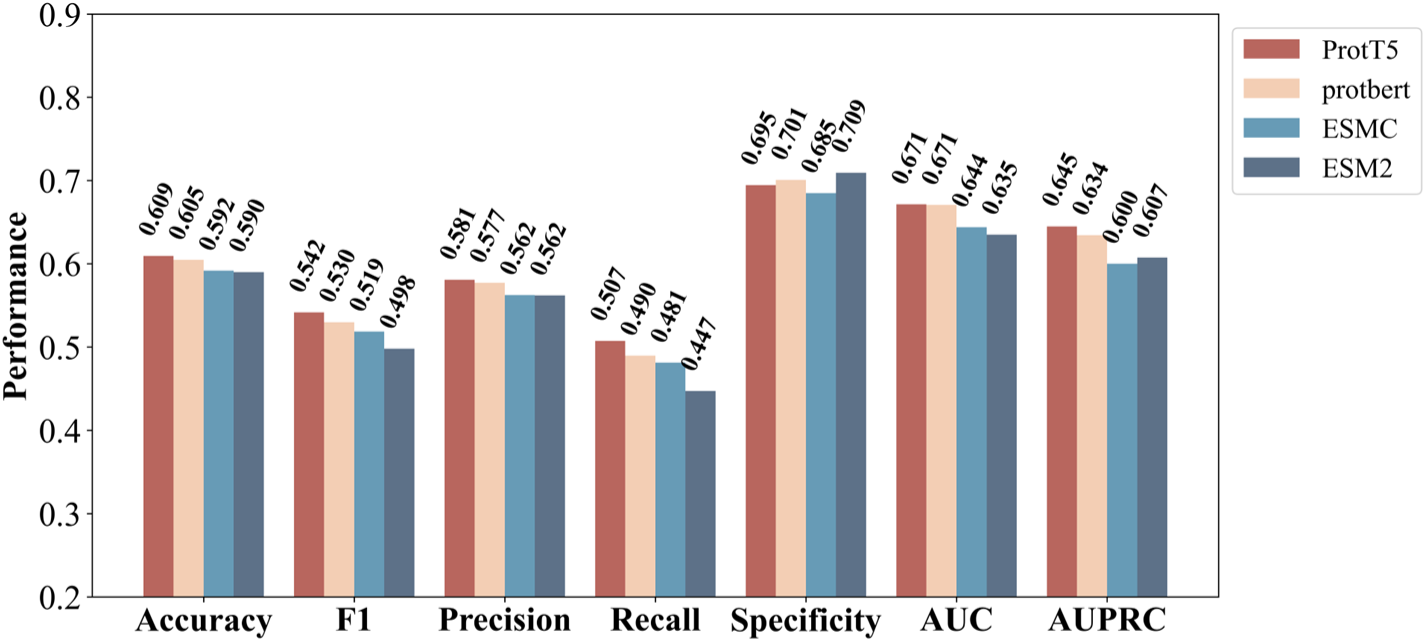
Performance Comparison of AlloPED-site Using Features from Different Protein Language Models.

After determining the optimal window size, we further compared the performance of features extracted from different protein language models in the task of predicting allosteric sites. The results indicated that the ProtT5 model performed the best, achieving an AUC of 0.671 and an AUPRC of 0.645, while also obtaining the highest MCC value of 0.205. Compared to other models, ProtT5 demonstrated a good balance in precision (0.580) and recall (0.507). The overall performance of the ProteinBERT model[32] was a close second to ProtT5, with an AUC of 0.670 and an AUPRC of 0.634; its precision reached 0.577, but its recall was relatively low at 0.490. Although the ESM series models[33, 34] performed well in terms of specificity, with ESM-2 reaching 0.709, their performance on other evaluation metrics was relatively weaker. The MCC metric was 0.162 for ESM-2, which has 650 million parameters, and 0.169 for ESM Cambrian (ESM C), which contains 600 million parameters. Despite their large scale, these models showed suboptimal predictive performance for allosteric sites. These results suggest that the ProtT5 model is more effective at capturing feature information related to allosteric sites within protein sequences, providing a more reliable feature representation for subsequent prediction tasks.

To validate the reliability of our model, we evaluated it using a test set of 24 allosteric proteins developed for the AllositePro method and compared the results with those of other widely used allosteric site identification methods on the same test set. By combining the pockets predicted by AlloPED-pocket, AlloPED-site further predicted allosteric sites, achieving a precision of 0.601, a recall of 0.4218, and a specificity of 0.661. Furthermore, to assess the impact of protein language model (PLM)-derived features, we incorporated sequence-based representations generated by ESM-2, ESM C, ProteinBERT, and ProtT5, then combined them with structural features for model evaluation (Table 2). The results indicate that the choice of PLM significantly influences predictive performance. Notably, ProtT5-based features yielded the best results, achieving a precision of 0.601, a recall of 0.422, and an AUC of 0.563, outperforming other PLM-derived feature combinations. In contrast, features derived from ESM-2 and ESM C exhibited lower recall values (0.360 and 0.378, respectively), indicating potential limitations in capturing relevant allosteric site information. These findings emphasize the effectiveness of ProtT5 in enhancing allosteric site prediction when integrated with structural descriptors, further demonstrating the advantage of combining sequence-based deep learning with structural insights.

## 4 Conclusions

This study introduces AlloPED, a novel framework integrating ensemble learning and deep neural networks to address the challenge of predicting allosteric sites in proteins. By combining AlloPED-pocket—an ensemble model optimized for pocket-level prediction using residue contact networks and physicochemical features—and AlloPED-site—a dilated convolutional neural network augmented with ProtT5 protein language model embeddings and attention mechanisms—our approach achieves state-of-the-art performance on benchmark datasets. AlloPED-pocket yields an MCC of 0.544 and AUC of 0.920, surpassing existing methods such as AllositePro (MCC=0.388) and PARS (MCC=0.099). AlloPED-site further refines predictions at the residue level, achieving a precision of 0.601 and recall of 0.422, demonstrating the value of multimodal feature integration.

The success of AlloPED underscores the importance of combining evolutionary and structural insights with advanced machine learning techniques. Key features such as residue clustering coefficients and van der Waals volume highlight the role of network connectivity and compact structural properties in allosteric regulation. This framework not only advances computational prediction of allosteric sites but also provides mechanistic insights into their functional roles, with implications for drug discovery targeting proteins like GPCRs and kinases. Future work will focus on integrating dynamic features from molecular dynamics simulations and validating predictions through experimental mutagenesis, further enhancing the framework’s utility in precision medicine and structural biology.

## Supplementary Materials

**Table S1.** The 19 physicochemical features provided by Fpocket.

| Feature Symbol | Description |
| --- | --- |
| fpocket_Score | Pocket score |
| fpocket_DruggabilityScore | Pocket druggability score |
| fpocket_NumberofAlphaSpheres | Number of Alpha Spheres in the pocket |
| fpocket_TotalSASA | Total SASA in the pocket |
| fpocket_PolarSASA | Polar SASA in the pocket |
| fpocket_ApolarSASA | Apolar SASA in the pocket |
| fpocket_Volume | Pocket volume |
| fpocket_Meanlocalhydrophobicdensity | Mean local hydrophobic density in the pocket |
| fpocket_Meanalphasphereradius | Mean alpha sphere radius in the pocket |
| fpocket_Meanalpsphsolventaccess | Mean alpsph solvent access in the pocket |
| fpocket_Apolaralphasphereproportion | Apolar alpha sphere proportion in the pocket |
| fpocket_Hydrophobicityscore | Pocket hydrophobicity score |
| fpocket_Volumescore | Pocket volume score |
| fpocket_Polarityscore | Pocket polarity score |
| fpocket_Chargescore | Pocket charge score |
| fpocket_Proportionofpolaratoms | Proportion of polar atoms in the pocket |
| fpocket_Alphaspheredensity | Alpha sphere density in the pocket |
| fpocket_CentofmassAlphaSpheremaxdist | Cent of mass alpha sphere max dist in the pocket |
| fpocket_Flexibility | Pocket flexibility |

**Table S2.**
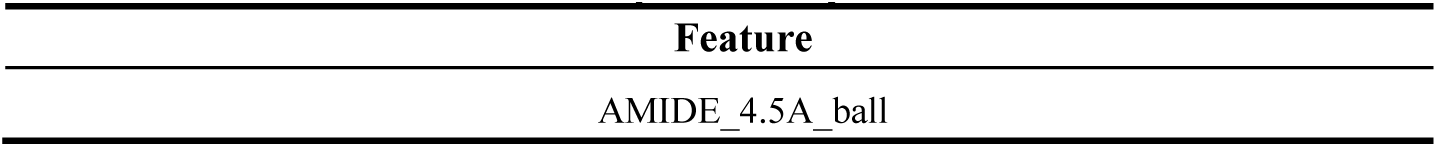
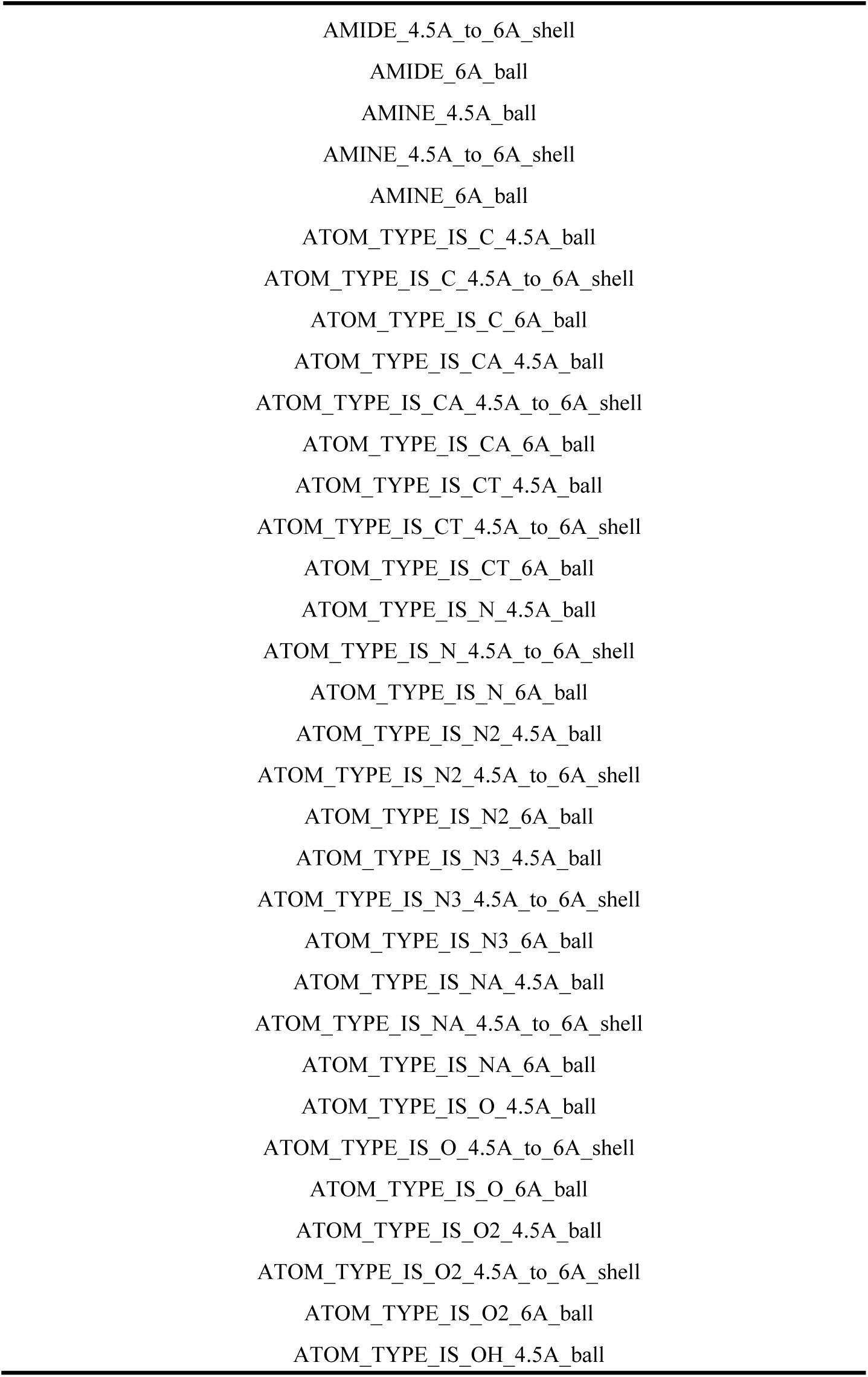
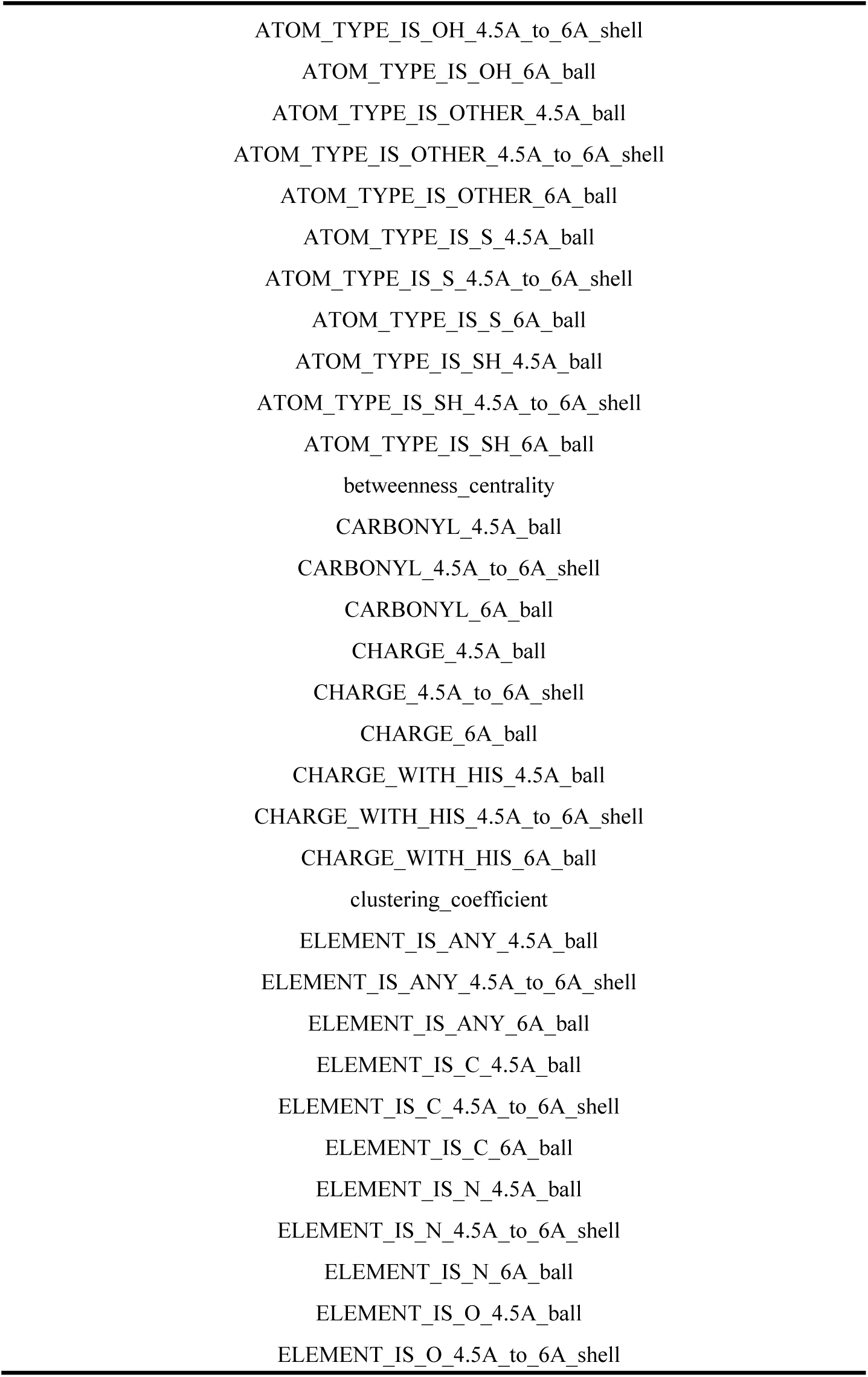
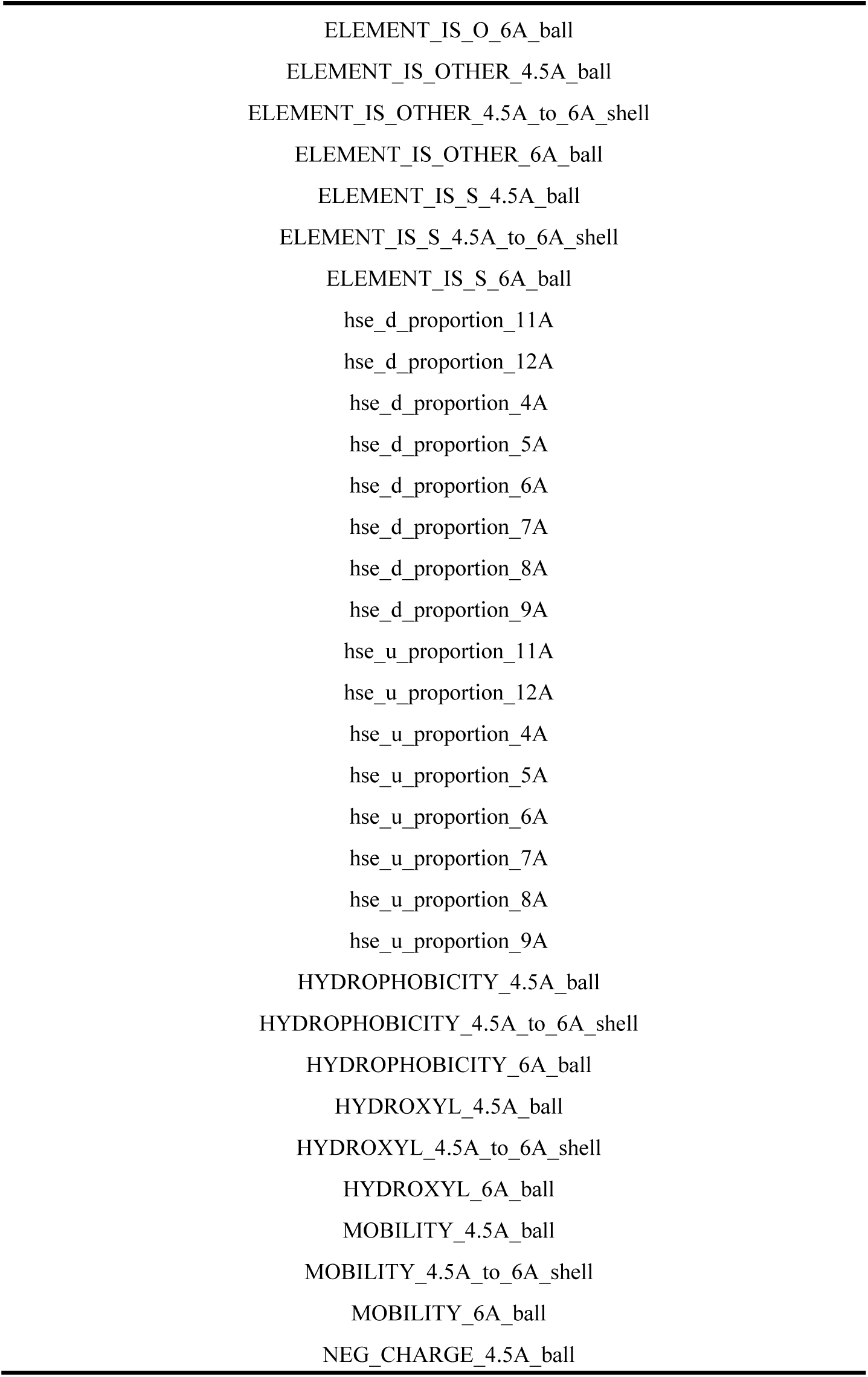
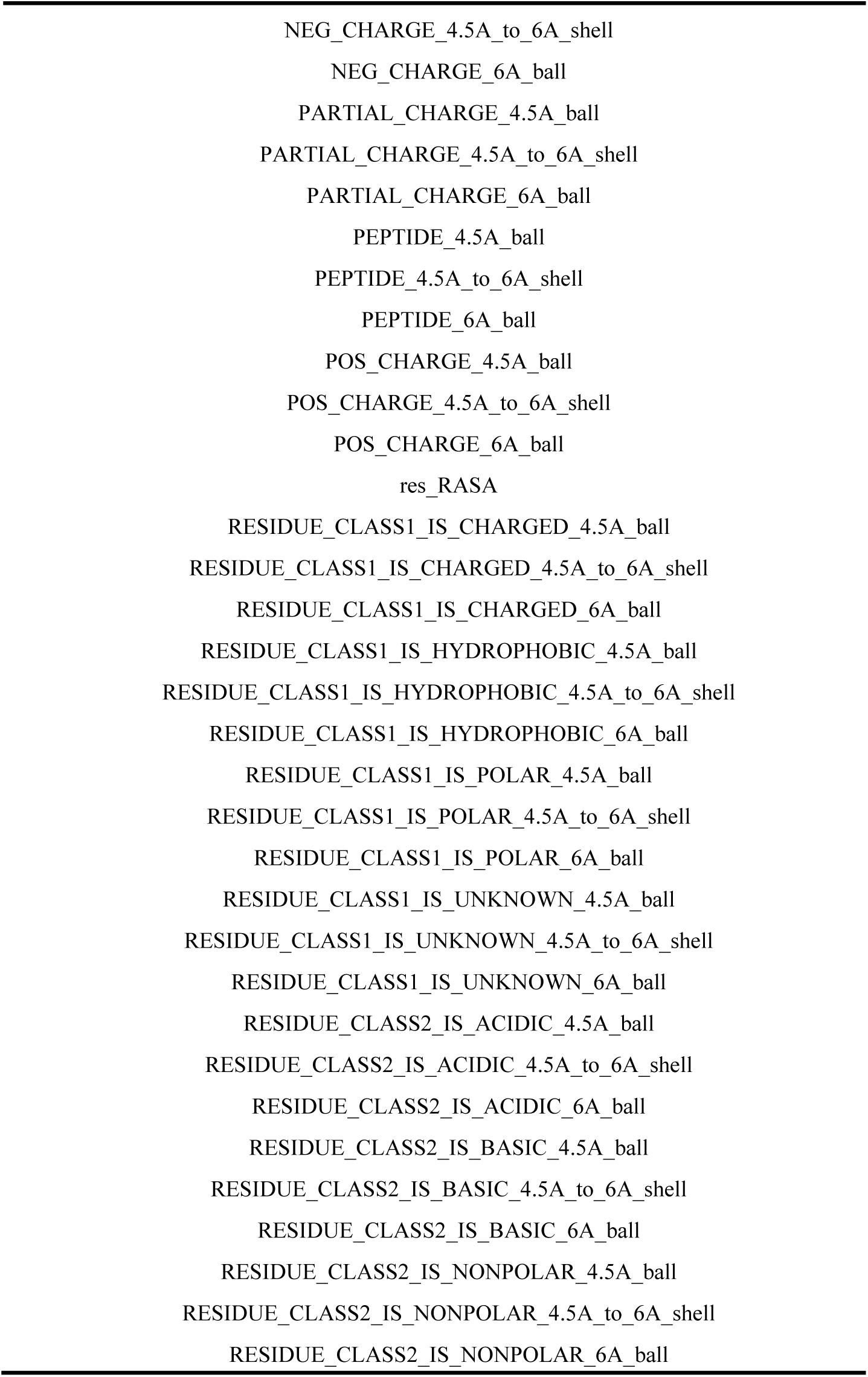
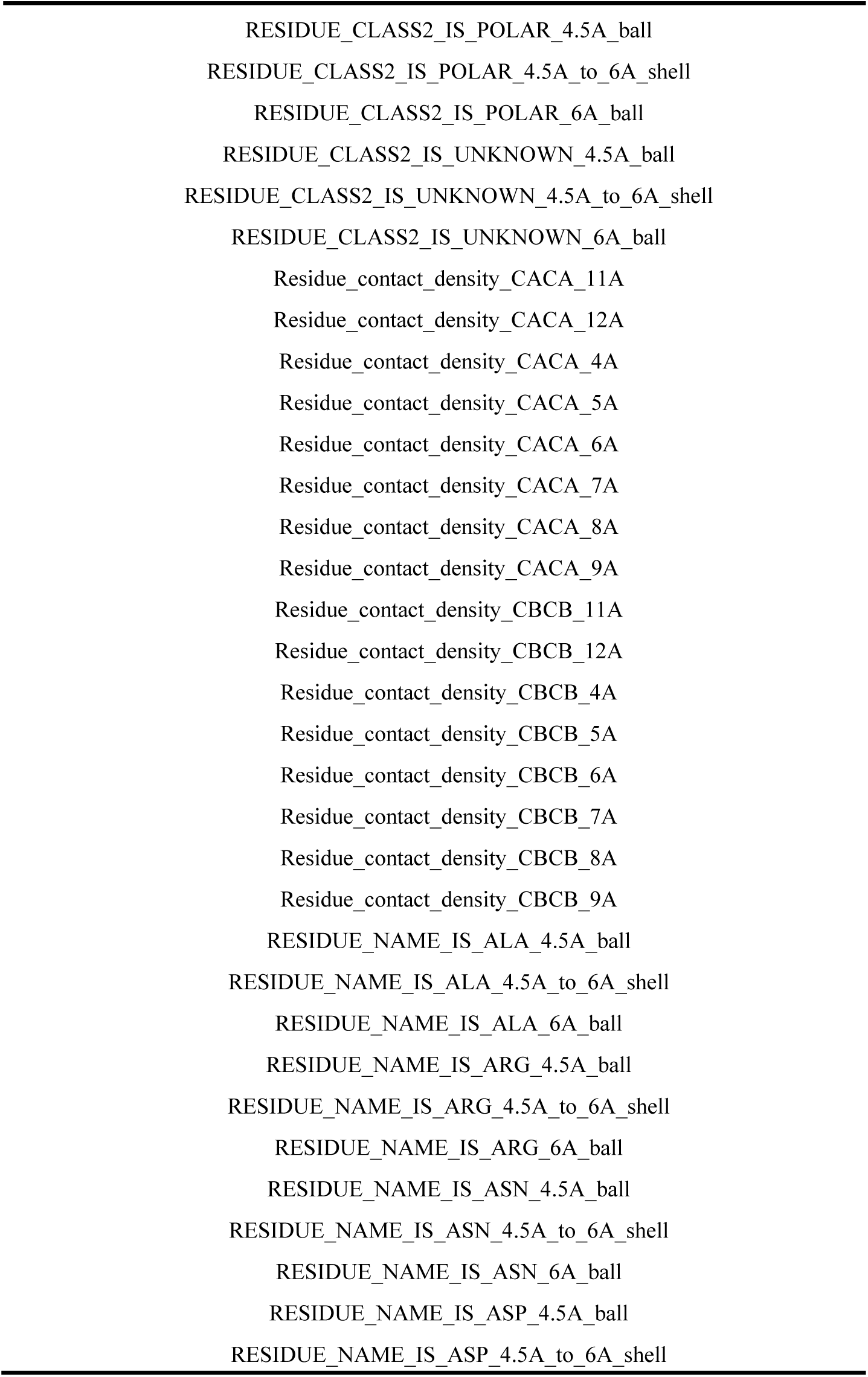
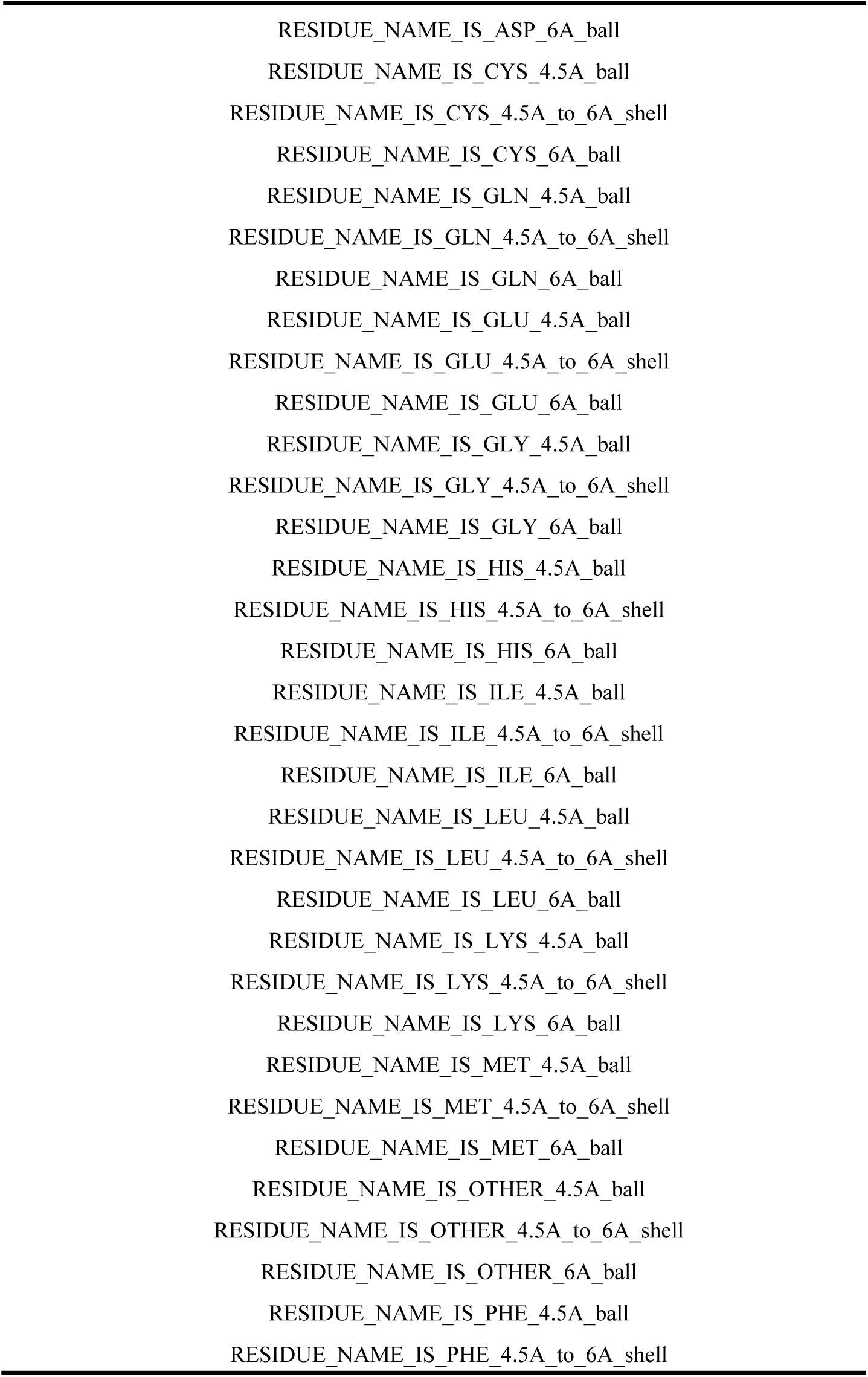
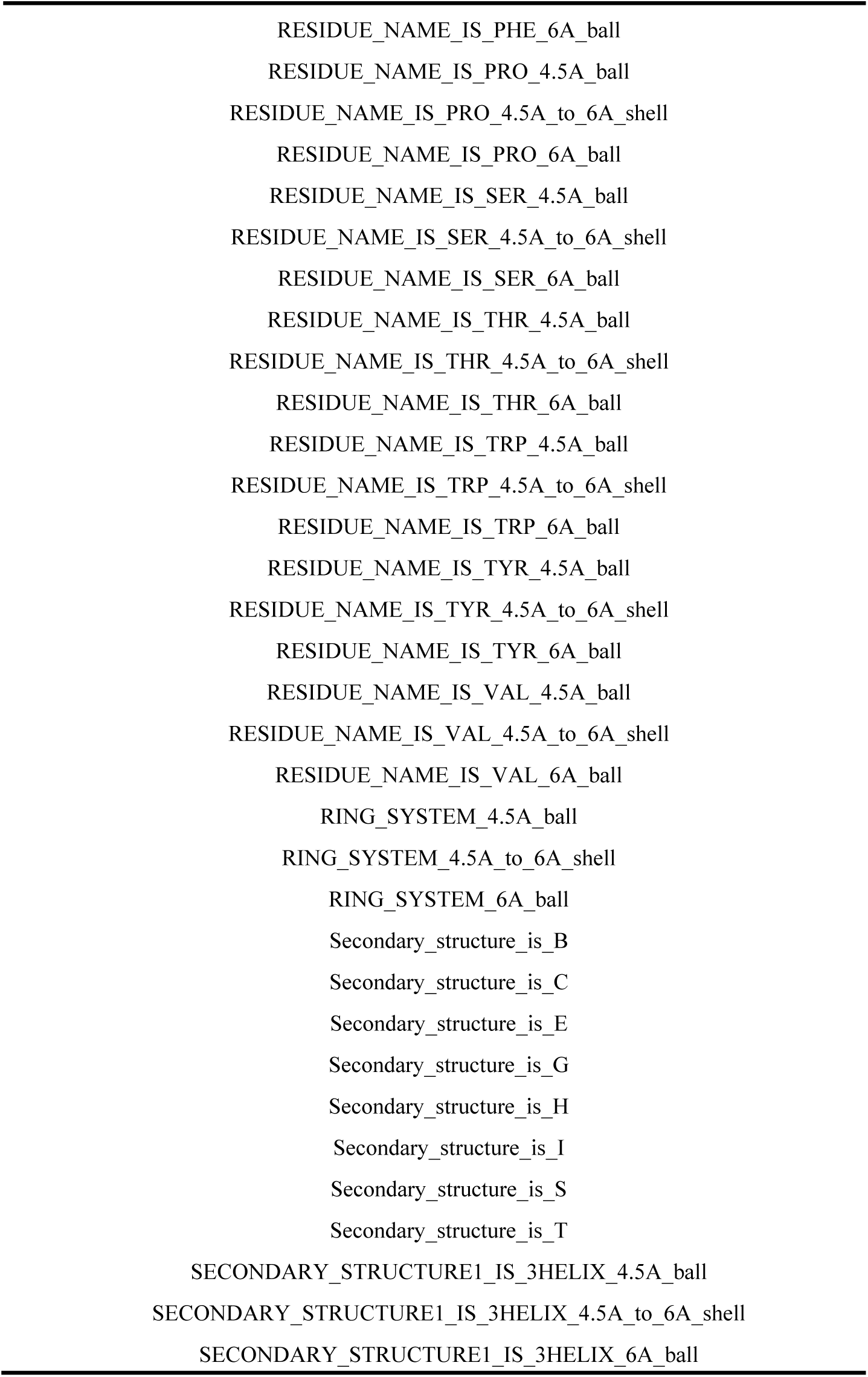
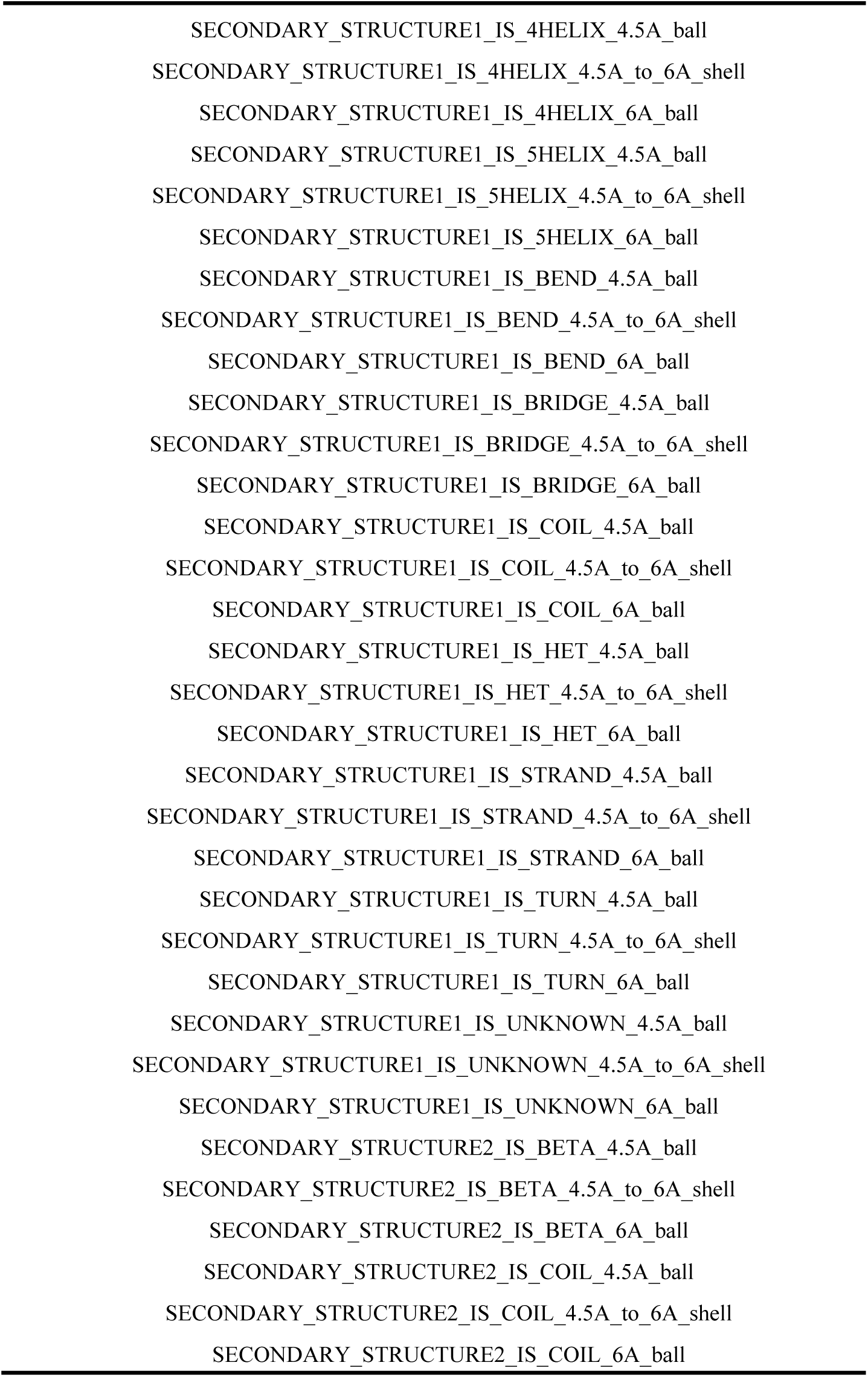
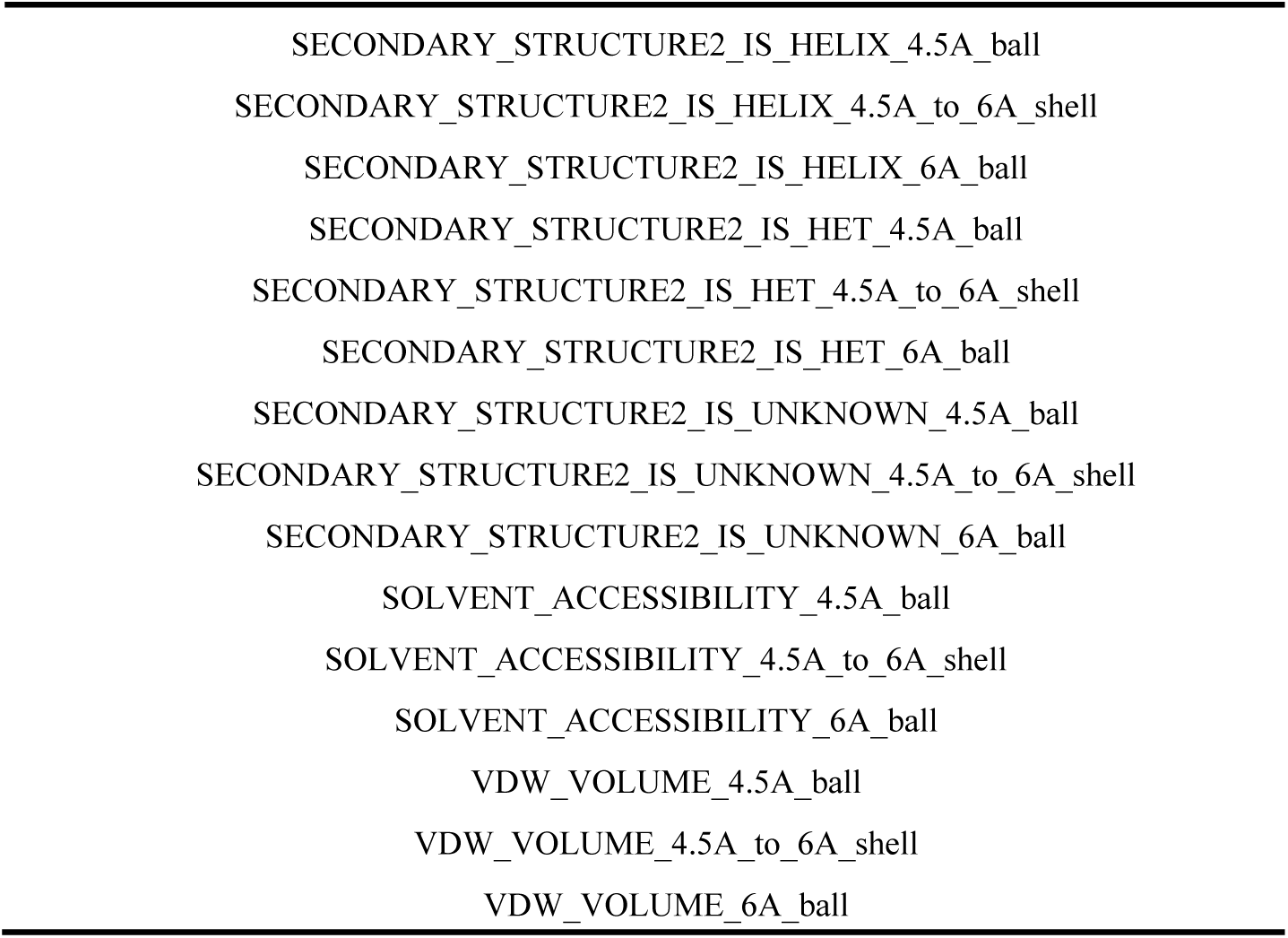
The structural features provided by PNFx.

## Notes

### Competing Interest Statement

The authors have declared no competing interest.

